# Single-Cell Analysis of Meningiomas Reveals Mutation-Associated Tumor and Immune Cell Gene Expression Programs

**DOI:** 10.1101/2025.05.21.655421

**Authors:** Jose A. Maldonado, Bryce L. Mashimo, Anthony Z. Wang, Rupen Desai, Saad M. Khan, Ngima D. Sherpa, Markus I. Anzaldua-Campos, Gregory J. Zipfel, Albert H. Kim, Joshua L. Dowling, Eric C. Leuthardt, Joshua W. Osbun, Ananth K. Vellimana, Michael R. Chicoine, Gavin P. Dunn, Allegra A. Petti

## Abstract

Genomic and epigenetic profiling, particularly DNA methylation analysis, have refined the molecular classification of meningiomas and revealed marked intratumoral heterogeneity. To further characterize heterogeneity within and between patients, we analyzed meningiomas using single-cell RNA sequencing (*n*=11), whole-exome sequencing (*n*=9), and spatial transcriptomics (*n*=3). Single-cell analysis revealed six transcriptionally distinct tumor cell states that corresponded to unique biological processes. Integration of the single-cell data with published exome and bulk RNA sequencing data from a large cohort revealed significant associations among somatic variants, tumor cell states, and immunological signatures. Notably, *NF2*-altered tumors were enriched for an epithelial-to-mesenchymal transition (EMT) cell state and immune cells, whereas *NF2*-intact tumors were enriched for a sterol metabolism cell state. Spatial transcriptomic analysis confirmed co-localization of immune cells and EMT tumor cells. Comparisons with immune cells from other brain tumors and peripheral tissues highlighted immunological cell states specific to meningioma. Collectively, these findings refine our genetic and molecular understanding of meningioma heterogeneity and underscore the link between genotype and molecular phenotype.

## Introduction

Meningiomas are the most common primary intracranial tumors, comprising more than one-third of all central nervous system neoplasms^1^. Meningiomas are classified into three WHO grades based on histopathological and molecular features. Grade I meningiomas are benign and account for about 80% of cases, with a 10-year recurrence rate of 7–25%. Grade II (atypical) meningiomas exhibit increased mitotic activity or brain invasion, and recur in 29–52% of cases within 5 years. Grade III (anaplastic or malignant) meningiomas are highly aggressive, often showing marked anaplasia and brain infiltration, with recurrence rates as high as 94% and significantly shortened survival. However, conventional histopathological subtyping alone has limited prognostic utility, particularly for low-grade meningiomas, underscoring the need for molecular characterization^2–6^. Recurrence risk is influenced by numerous additional variables, including completeness of surgical resection, specific genetic alterations, and tumor location (which is strongly associated with genetic alterations^7^). In particular, genetic heterogeneity is increasingly recognized as central to the biological behavior and clinical management of meningiomas.

Early large-scale genomic studies identified mutations in *NF2*, *TRAF7*, *KLF4*, *AKT1*, and *SMO* as early driver mutations in meningioma and established the initial framework for stratification of meningioma into genomic subtypes^8–10^. These genetic alterations correlate with clinical, histological, and anatomical subtypes: *NF2*-mutant meningiomas are typically fibrous in histology and originate from the convexity, whereas non-*NF2*–mutant tumors more commonly arise from the skull base and display a broader range of histopathological features^11,12^. Tumor genetics also correlates with WHO grade, such that higher-grade meningiomas tend to carry a higher mutation burden and more complex genomic profiles^13^. The 2021 WHO classification of CNS tumors recognized the significance of genetic profiling by incorporating molecular features into the classification framework for meningiomas^14^.

To complement genomic studies, DNA methylation profiling has been used to stratify meningiomas into clinically meaningful groups, but these do not map cleanly to the WHO grades. In particular, DNA methylation has demonstrated that even tumors of the same grade can belong to different epigenetic subgroups, with some histologically benign tumors exhibiting methylation patterns characteristic of aggressive meningiomas^15^. More recently, integrative molecular approaches have been used to suggest alternative classifications and elucidate meningioma biology. These studies have combined genome sequencing, RNA sequencing, DNA methylation profiling, and cytogenetics to identify new molecular subtypes that better correlate with prognosis and suggest new therapeutic vulnerabilities^8,9,15–17^. In addition, several studies have used single-cell RNA-sequencing (scRNA-seq) to describe intratumoral cellular and spatial heterogeneity within meningiomas which may contribute to ambiguity in molecular classification^18–20^. As such, relationships among tumor genetics, intratumoral heterogeneity, and the various histological and molecular classifications are not well understood.

To better understand the relationships among WHO grade, tumor genetics, tumor cell state, and the immune microenvironment in meningioma, we integrated single-cell RNA sequencing (scRNA-seq) and whole-exome sequencing across a cohort of primary sporadic meningiomas. We identified six transcriptionally distinct tumor cell states reflecting diverse biological processes, such as epithelial-mesenchymal transition (EMT), sterol metabolism, and chromatin remodeling. Using published exome and bulk RNA-sequencing data from a larger cohort^13^, we found that these tumor cell states are correlated with specific genotypes—such as *NF2* alterations and *TRAF7*/*KLF4* mutations—and with the composition of the immune microenvironment. In particular, *NF2*-mutated tumors were enriched for EMT tumor cells and *CD8+* T-cells, whereas *NF2*-intact tumors were enriched for sterol-metabolic tumor cells. We performed spatial transcriptomics on a subset of these samples, which confirmed co-localization of EMT tumor cells with *CD8*⁺ T cells and suggested mechanisms of immunosuppression in meningioma. Ligand-receptor analysis revealed distinct patterns of intercellular communication between the immune cells and the different tumor cell states. Finally, we used scRNA-seq to compare immune cells across multiple tissues (meningioma, glioblastoma, vestibular schwannoma, nonmalignant dura, and peripheral blood), which demonstrated that meningioma-associated immune cells exhibit unique transcriptional programs. Collectively, our study connects genetic heterogeneity, transcriptional heterogeneity, and composition of the immune microenvironment in the context of meningioma.

## Results

### Heterogeneous genomic landscape of meningioma tumors

To characterize the cellular and molecular heterogeneity of human meningiomas, we performed single-cell RNA-sequencing (scRNA-seq, *n* = 11), whole exome sequencing (WES, *n* = 9), and spatial transcriptomics (*n* = 3) on freshly resected primary sporadic meningiomas (Fig. 1A). These tumors represented diverse clinical and pathological features, including variation in age, sex, tumor size and location, histological grade, and genetic alterations (Fig. 1B, C, Supp. Table 1).

**Fig 1.**
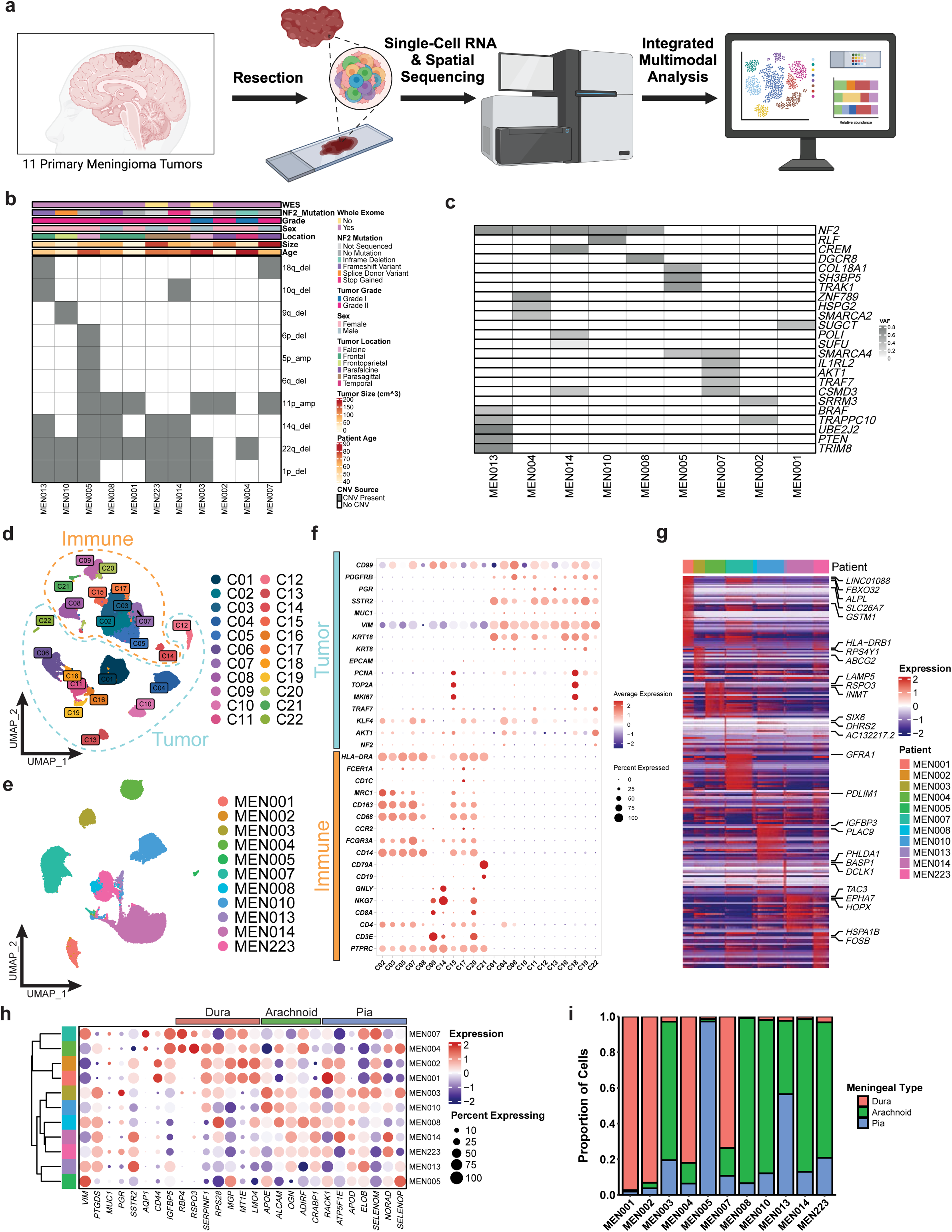
Genomic profiling of meningioma tumors. **a,** Schematic of the study workflow. Tumor samples from 11 primary meningiomas underwent resection, single-cell RNA and spatial sequencing, followed by integrated multimodal analysis. **b,** Heatmap displaying chromosomal deletions across samples, with annotations for key clinical and molecular features. **c,** Mutation profile of meningiomas illustrating key mutations across samples, with shaded boxes indicating detected mutations. **d,** Uniform manifold approximation and projection (UMAP) of single-cell transcriptomes; Clustering of immune (orange) and tumor (blue) cell populations, with identified clusters labeled. **e,** UMAP plot highlighting inter-patient heterogeneity in sc-RNAseq data. **f,** Dot plot showing expression levels (color) and percent of cells expressing key genes across meningioma samples from sc-RNAseq data. **g,** Heatmap of differentially expressed genes; Hierarchical clustering of gene expression across samples, with expression levels scaled from low (blue) to high (red). **h,** Dot plot displaying meningeal layer-specific gene expression patterns across individual patient tumor cells. **i,** Bar plot showing the proportion of predicted meningeal layer similarity for individual patients’ meningioma tumor cells.

To characterize the genomic landscape of this cohort, we used WES to identify somatic variants, and a combination of scRNA-seq and WES to identify copy-number variants (CNVs). From this we observed SNVs ranging between 1 to 25 per patient, with a median of 12 identified (Supp. Table 2). Previous large-scale studies of meningioma^3,16,21^ established two main genomic subtypes defined by mutually exclusive alterations in *NF2* (which can occur through mutation and/or deletion within Chromosome 22^22^) and *TRAF7*. Our cohort reflected this dichotomy, such that 9 of 11 tumors contained an *NF2*-spanning deletion in Chr22, and five of these (MEN008, MEN013, MEN014, MEN010, and MEN004) also contained single nucleotide variants (SNVs) in *NF2* (Supp. Table 3). Of the remaining two patients with WES data, one case (MEN007) harbored *TRAF7* and *AKT1* mutations. The other (MEN002) did not belong to either category, and contained SNVs in genes such as *TRAPPC10* and *SRRM3* (Fig. 1C). Our cohort also exhibited a variety of other CNVs; multiple tumors contained concurrent deletions, the most common of which was Chr22q and Chr1p loss (6 tumors), and five tumors contained three or more concurrent CNVs (Fig. 1B, Supp. Table 3).

Next, we used the scRNA-seq data to analyze the cellular composition of the meningioma ecosystem. Following the removal of doublets and low-quality cells and correction for ambient RNA, our scRNA-seq dataset contained 24,638 genes and 84,130 cells (Extended Data Fig. 1A). Dimensionality reduction and graph-based clustering delineated putative populations of tumor and immune cells (Fig. 1D). Subsequent analyses based on CNVs and marker genes identified 38,397 tumor cells and 45,733 immune cells (Fig. 1D-F). Briefly, each cluster was further characterized based on the expression of established marker genes for immune cells (e.g. *PTPRC/CD45, CD3E, CD14, CD68, MRC1* and *CDC1)* and meningioma cells (*VIM, SSTR2, PGR* and *PDGFRB)* (Fig. 1F, Supp. Table 4). Despite similarities in immune composition across tumors, the tumor cells exhibited notable interpatient heterogeneity (Extended Data Fig. 1B). To further explore these differences, we identified differentially expressed genes (DEGs) that distinguished each patients’ tumor cells (Fig. 1G; Supp. Table 5, 6, Extended Data Fig. 1C), and found that several DEGs recurred across multiple patients. For instance, the gene for alkaline phosphatase, *ALPL,* was strongly upregulated in MEN001, MEN002, and MEN007, with no expression in other tumors. Other DEGs shared by multiple patients included the matrix metalloproteinase inhibitor *TIMP3*, the cytochrome P450 gene *CYP1B1*, ABC transporter *ABCG2*, Y-linked ribosomal protein *RPS4Y1*, which, as expected, was expressed solely in male patients, long non-coding RNA *LINC01088,* WNT regulator *SFRP4* insulin-like growth factor binding protein *IGFBP5*, and F-box protein *FBXO32. IGFBP5* is a marker of immune infiltration and poor prognosis in glioma; it was also identified as a highly connected component of an oncogenic module in a meningioma network analysis based on microarray data^23,24^. Interestingly, *PTGDS* was highly expressed in nearly all MEN003 and MEN014 cells, exhibited bimodal expression in MEN004, MEN008, MEN010, MEN013, and MEN223, and displayed minimal expression in MEN001, MEN002, MEN005 and MEN007 (Fig. 1G; Extended Data Fig. 1A, Supp. Tables 5, 6, Extended Data Fig. 1C).

Previous studies suggest that meningiomas can arise from cells of the meningeal layers including dural border cells and arachnoid cells that express PGDS, the protein encoded by the *PTGDS* gene^25,26^. Given this relationship, we characterized our samples using published gene expression data from fibroblasts derived from each meningeal layer^27^. This analysis showed that marker genes for the dura, arachnoid, and pia mater layers displayed distinct, patient-specific expression patterns (Fig. 1H, Extended Data Fig. 1D). When we scored the cells in each tumor for composite expression of these layer-specific genes, we found that each tumor was primarily composed of cells derived from a single layer (Supp. Table 7, Extended Data Fig. 1E). MEN001, MEN002, MEN004, and MEN007 scored most highly for the dura mater expression signature, while MEN005 scored most highly for the pia mater signature, and the remaining tumors expressed the arachnoid mater signature (Fig. 1I). Taken together, our exome and scRNA-seq data corroborated known genetic subtypes, highlighted genetic and transcriptional differences between patients, and suggested potential cell-of-origin signatures.

### Meningiomas are composed of six tumor cell states that exhibit unique gene expression programs and biological functions

To characterize intratumoral heterogeneity in meningioma, we used non-negative matrix factorization (NMF) to identify the tumor cell states that comprise individual meningiomas and are conserved across patients. We iterated over a range of ‘k’ values and filtering parameters, which resulted in 50 gene expression programs. Hierarchical clustering of these programs revealed six gene expression “modules” that were shared across patients and robust to parameter selection (Figure 2A; Supp. Table 8, 9). These modules were: “Cell Cycling” (M1), “Chromatin Remodeling” (M2), “Sterol Metabolism” (M3), “EMT (Epithelial-Mesenchymal Transition)” (M4), “Stress Response” (M5), and Hypoxia (M6). We then used the Toppfun and Cytoscape tools to characterize the functional enrichment of the genes defining each module (Supp. Table 8, 9). Each module was enriched for specific biological processes (Fig. 2B, Supp. Table 8). Module 1 (M1), the smallest module (4.1% of cells), was enriched for cell cycle genes (*STMN1, TYMS, TOP2A*). Functional enrichment showed associations with mitotic activity through upregulation of chromosome segregation genes, and a predominance of cells in the G2/M phase. In our single-cell data, the Cell Cycling (M1) module was most represented in higher grade tumors and this association was also recapitulated in external meningioma scRNA-seq datasets^19,28^. Module 2 (M2) (20% of cells) was enriched for chromatin remodeling genes (*SETD2, ARID1B, ASH1L*) and long non-coding RNAs (*SNHG14, KCNQ1OT1*) implicated in tumorigenesis^29^. MEN014, highly represented in M2, harbored mutations in *POLI* and *CSMD3*, genes linked to high tumor mutational burden^30^. Module 3 (M3) (23.3% of cells) featured sterol metabolism genes (*FADS1, PTGES, SCD*) and was enriched in cholesterol biosynthesis pathways. Our single-cell dataset revealed an enrichment of this cell module in patients with Chr11p-amplified tumors, a chromosomal region harboring genes previously linked to dysregulated lipid metabolism and lipid-associated pathologies^31,32^. Module 4 (M4) (28.5% of cells), the most abundant module, was enriched for ECM remodeling and vascular-related genes (*COL1A1, LAMA2, VCAM1*), consistent with meningioma-associated NF2 mutations affecting ECM regulation and epithelial-to-mesenchymal (EMT) transition^33^. Module 5 (M5) (11.5% of cells) was defined by apoptotic and AP-1, or stress response, transcription factor targets (*BCL3, FOS, JUNB*), and was enriched for biological processes associated with hormonal activity, including glucocorticoid signaling. Module 6 (M6) was enriched for HIF-1α-regulated apoptotic genes (*IGF2, H19*); *IGF2* is implicated in tumor proliferation and hypoxic adaptation.

**Fig 2.**
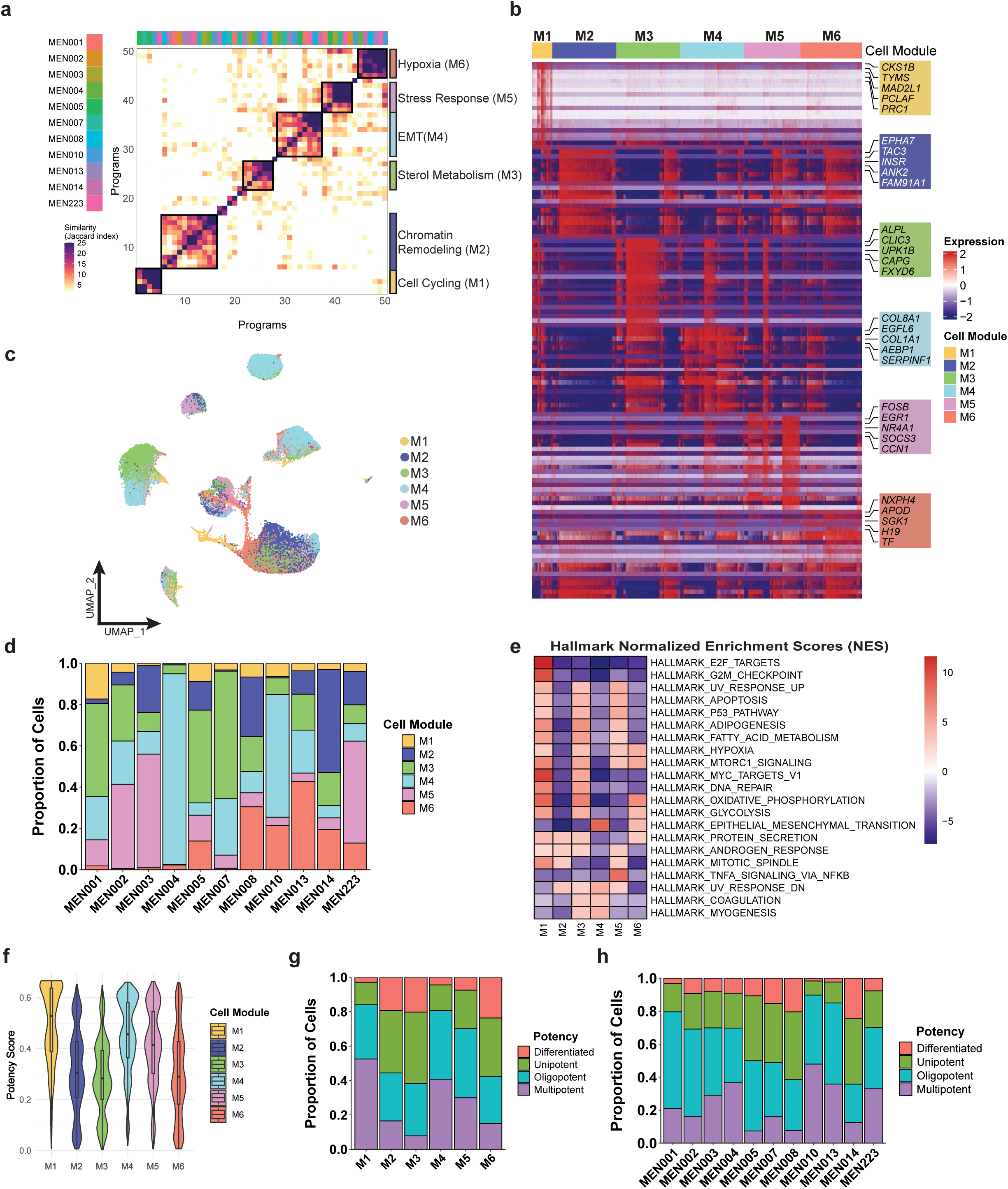
NMF cell module characterization. **a,** Clustered heatmap of transcriptional programs derived from NMF across meningioma samples, highlighting major biological processes associated with each module. **b,** Heatmap of differentially expressed genes across transcriptional modules, with the top 5 marker genes highlighted. **c,** UMAP visualization of single-cell transcriptomes, with cells colored by assigned transcriptional module. **d,** Bar plot showing the proportion of cells assigned to each transcriptional module across meningioma tumors from individual patients. **e,** Hallmark gene set enrichment analysis displaying normalized enrichment scores (NES) for each transcriptional module. **f,** Violin plot of potency scores as across transcriptional modules, indicating variation in cellular states. **g,** Bar plot showing the proportion of cells classified by potency across transcriptional modules. **h,** Bar plot displaying potency classification across individual patients’ meningioma tumors.

To confirm these findings in an independent scRNA-seq data set, we expanded the data set to include five meningioma samples from Choudhury et. al and one Grade III sample from Wang et. al^16,28^. We repeated the NMF analysis and identified similar modules shared across patients (Extended Data Fig. 2A), recapitulating our findings (Extended Data Fig. 2B, p < 0.05, hypergeometric test). We refer to these modules as the Cycling (M1), Chromatin (M2), Sterol (M3), EMT (M4), Stress (M5), and Hypoxia (M6) modules. Together, these modules provide insight into the functionally specialized tumor cell populations that comprise each meningioma.

Next, we interpreted each gene module as a tumor “cell state”, and assigned each tumor cell to one of the six cell states by determining which gene module was most highly expressed in each cell (Fig. 2C, Methods). The results revealed both intratumoral and intertumoral heterogeneity: although every cell state was present in every patient, each tumor was composed of varying proportions of each cell state which reflected the unique composition of each tumor (Extended Data Fig. 2C). Some cell states comprised as much as 92.5% of a tumor (EMT (M4) in MEN004) or as little as 0.14% (Stress (M5) in MEN004)(Fig. 2D, Supp. Table 10).

To characterize meningioma tumor cell states more deeply, we used the fGSEA R package to assess the relative expression of the Hallmark, KEGG, and Gene Ontology pathways in cells assigned to each cell state. This indicated that different tumor cell states utilized different pathways (Fig. 2E, Supp. Table 11, Extended Data Fig. 2D, E). Specifically, Cycling (M1) tumor cells exhibited upregulation of the E2F targets (11.63), MYC targets v1 (10.57), and G2M checkpoint (10.05) Hallmark pathways. Chromatin (M2) cells, on the other hand, exhibited upregulation of Hallmark pathways related to protein secretion (3.27), and downregulation of UV response pathways (3.26) and the mitotic spindle (3.09). In Sterol (M3) cells, the top three upregulated Hallmark pathways were oxidative phosphorylation (7.11), adipogenesis (6.27), and fatty acid metabolism (5.57), suggesting a high level of metabolic activity in this cell state. EMT (M4) cells, on the other hand, exhibited upregulation of Hallmark pathways related to the epithelial-to-mesenchymal transition (8.73), angiogenesis (2.17), and inflammatory response (1.56), suggesting modulation by diverse immune and non-immune cells in the tumor microenvironment. Stress (M5) cells reflected stress response mechanisms, with the top upregulated Hallmark pathways including TNFA signaling via NFKB (8.11), the P53 pathway (4.89), and UV response (4.73). Lastly, Hypoxia (M6) cells exhibited upregulation of oxidative phosphorylation (6.51), MTORC1 signaling (5.07), and hypoxia (4.63), reflecting the utilization of multiple oxygen-related pathways.

To understand whether different cell states are associated with different levels of differentiation or stemness, we assessed their “developmental potency” using CytoTRACE2. “Potency” was assessed by analyzing the transcriptional diversity and expression patterns of genes associated with stemness derived from multiple tissues, rather than a predefined set of stemness genes^34^. The six cell states separated into two distinct groups based on potency scores: Chromatin (M2), Sterol (M3), and Hypoxia (M6) cells exhibited higher levels of differentiation, while Cycling (M1), EMT (M4), and Stress (M5) cells were more stem-like (Fig. 2F). Projecting these cell state annotations onto each tumor revealed substantial differences in the ratio of multipotent cells to differentiated cells. For example, MEN010 contained 47.96% multipotent cells and 1.67% differentiated cells, whereas MEN014 contained 12.65% multipotent cells and 24.33% differentiated cells (Fig. 2H, Supp. Table 12). Collectively, our results revealed six distinct, functionally specialized tumor cell states in meningiomas, each defined by unique transcriptional programs. To augment these findings, we then investigated how these states correlated with tumor genotypes and their surrounding immune microenvironment.

### Tumor cell states are correlated with tumor genetics and the tumor-immune microenvironment

Because each tumor has a unique distribution of tumor cell states, we hypothesized that cell state composition is associated with other molecular and clinical variables. For increased statistical power, we leveraged published bulk RNA-seq data^13^ from 145 primary meningiomas that were thoroughly annotated with respect to age, sex, histological grade (WHO), recurrence, and genotype (Figure 3A, Supp. Table 13). To identify relationships among cellular composition, clinical parameters, and somatic variants, we first used CIBERSORTx to infer the composition of each tumor with respect to the single-cell states we identified from our data. The presence of these cell states was verified through module scoring of the bulk RNA-seq data and correlated with cell state composition. (Extended Data Fig. 3A).

**Fig 3.**
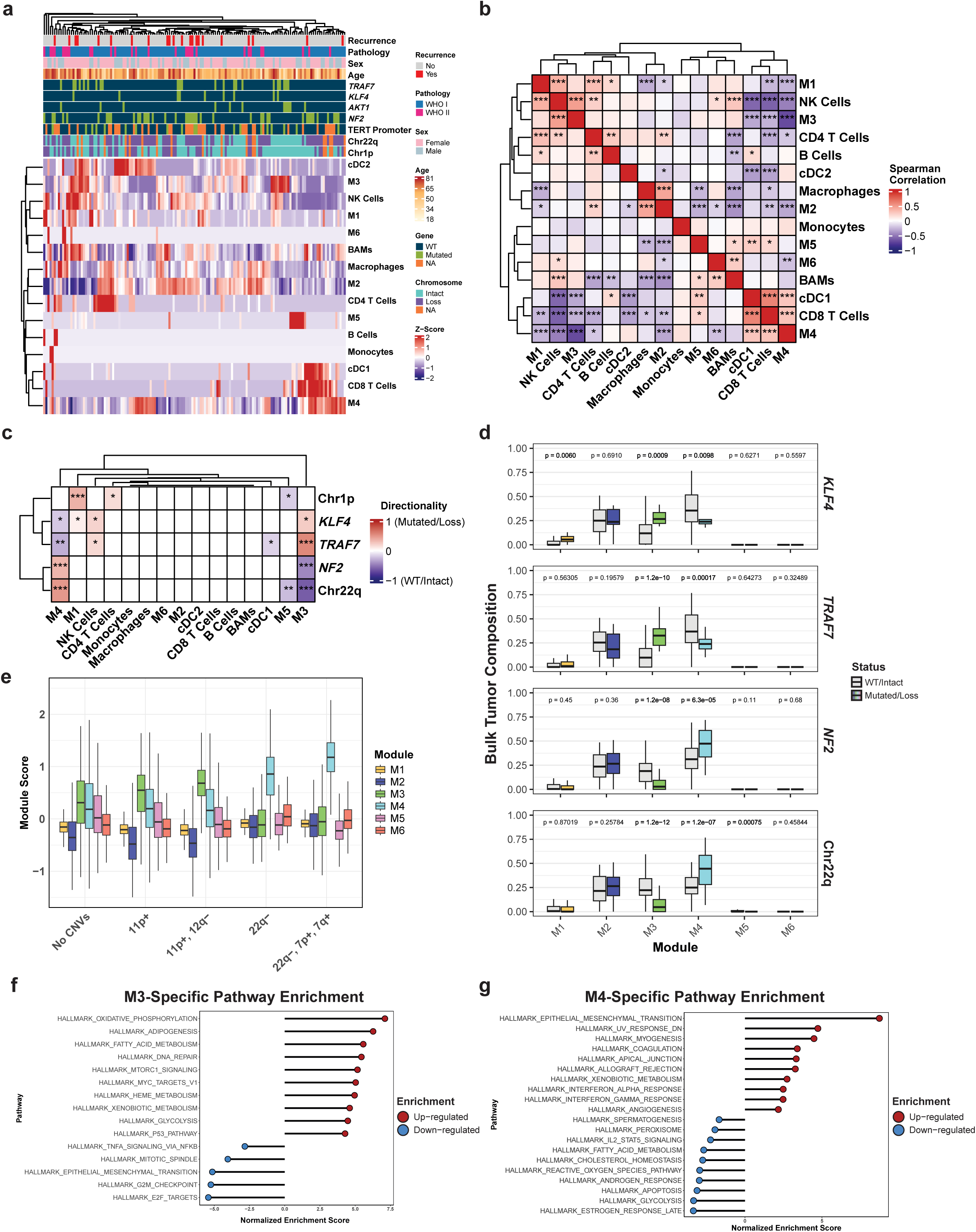
Bulk RNA-seq correlations. **a,** Heatmap of bulk RNA-seq deconvolution of tumor cell module and immune cell composition across meningioma samples, annotated with clinical and genomic features. **b,** Spearman correlation matrix heatmap displaying associations between transcriptional modules and immune cell populations. **c,** Heatmap of directional associations between genetic alterations and cellular composition. **d,** Box plots showing the cell module composition of meningioma tumors in relation to their *KLF4*, *TRAF7*, *NF2*, and Chr22q status. **e,** Module score distributions across different genetic alteration groups of tumor cells from scRNA-seq data. **f,** Pathway enrichment analysis for the M3 module, highlighting significantly enriched hallmark pathways. **g,** Pathway enrichment analysis for the M4 module, identifying key biological processes associated with this module.

We then examined relationships between tumor cell state and immune cell composition. We observed statistically significant correlations (*p* < 0.05, Spearman test) among 1) EMT (M4) tumor cells, cDC1s, and CD8 T Cells, 2) Chromatin (M2) tumor cells and macrophages, and 3) Cycling (M1) tumor cells, Sterol (M3) tumor cells, and NK cells (Fig. 3A). In addition, Hypoxia (M6) tumor cell abundance was correlated with border associated macrophage (BAM) abundance, but not significantly (Fig. 2B). While significant correlations between cell types and tumor cell states across patients were more pronounced and greater in number, fewer were identified between cellular composition or mutations and the patients’ clinical features such as age, sex, histological grade and recurrence status (Extended Data Fig. 3B-C).

Interestingly, we noted a marked anticorrelation between Sterol (M3) and EMT (M4) tumor cell states. To investigate this further, we then analyzed associations between genotype and tumor cell state composition using both the published exome data and our own scRNA-seq data. Importantly, the published cohort captured the dichotomy between *NF2*-altered tumors and *KLF4/TRAF7*-mutated tumors (Extended Data Fig. 3D). We found that the *NF2*-altered tumors were enriched for the EMT (M4) tumor cell state, while *KLF4/TRAF7*-mutated tumors were enriched for the Sterol (M3) tumor cell state (Fig. 3C-D). Thus, tumor genotype appears to influence tumor cell state composition. In our scRNA-seq data, cells without CNVs scored highly for the Sterol (M3) and EMT (M4) tumor states. However, those with Chr11p amplifications, either in isolation or in combination with other CNVs, consistently scored the highest for the Sterol (M3) cell module. Conversely, those with Chr22q losses score highly for the EMT (M4) cell module (Fig. 3E). The inverse relationship between the Sterol (M3) and EMT (M4) cell states is further highlighted through their functional enrichment. The pathways that are up-regulated and define the Sterol (M3) state, such as oxidative phosphorylation, fatty acid metabolism, and adipogenesis, are conversely downregulated in the EMT (M4) cell state. The inverse is also true with the epithelial to mesenchymal transition pathway being down-regulated in Sterol (M3) cells while being up-regulated and defining the EMT (M4) cell state (Fig. 3F-G, Supp. Table 14). Together, these results illustrate a direct link between tumor genotype, transcriptional cell state, and immune composition. The relationship was most stark in the dichotomy between Sterol (M3) and EMT (M4) phenotypes.

To evaluate the robustness of the M4 module and its association with Chr22q deletions in other tumor types, we scored tumor and immune cells from a vestibular schwannoma (VS)^35^ scRNA-seq data set with respect to our meningioma modules. In VS, the M4 module score was most pronounced in non-myelinating Schwann cells, known for their association with Chr22q deletion (Extended Data Fig. 3E). This suggests that the association between Chr22q deletions and the EMT (M4) cell state exists in multiple tumor types, and is pronounced in Chr22q loss or *NF2*-driven malignancies.

### Spatial transcriptomics confirms module-specific correlations

Next, we leveraged spatial transcriptomics to determine whether these cell state specific correlations are maintained *in situ*. To characterize the spatial organization of tumor and immune cells in a subset of samples, we performed spatial transcriptomics using the Visium technology on one sample from MEN006 and two samples from MEN007 (Extended Data Fig. 4A). After filtering for numbers of counts and features this yielded 1281, 2429 and 3414 high quality spots, respectively. Clustering of these spots (based on gene expression) yielded 8, 9 and 10 spatial clusters per sample (Fig. 4A, Supp. Table 15). Given the technical limitations of the Visium system, including its lack of single-cell resolution and inability to definitively assign transcriptomic profiles to individual cells, we then used the CellTrek R package to map cells from our scRNA-seq data to their most likely location in the tissue slices. This revealed that the spatial samples were composed primarily of tumor cells with a uniform, albeit lesser, presence of myeloid and lymphoid cells (Fig. 4B). CellTrek mapping enabled us to infer the cell type composition of each gene expression cluster (Extended Data Fig. 4B, Supp. Table 16). We then pooled spatial clusters across all samples, and clustered them based on their cell type composition (Fig. 4C). This showed that the EMT (M4) cell state is consistently localized with immune cells, particularly CD8 T cells, across all samples (Fig. 4C, Supp. Table 17), corroborating our sample-specific colocalization results as well as the bulk RNA-seq deconvolution analysis (Fig. 4D, Extended Data Fig. 4C). This relationship was also captured by CellTrek’s co-localization metrics (Extended Data Fig. 4D). To further investigate this, spatial clusters were stratified based on the abundance of EMT (M4) cells: clusters with a z-score > 0.5 were classified as M4-enriched, and those with a z-score < -0.5 as M4-depleted. Gene set enrichment analysis revealed that M4-enriched regions exhibited increased activity of epithelial-to-mesenchymal transition pathways and decreased activity in UV response and interferon gamma signaling. In contrast, M4-depleted regions showed upregulation of estrogen response pathways (early and late), E2F targets, and G2/M checkpoint pathways (Fig. 4E–F; Supp. Table 18).

**Fig 4.**
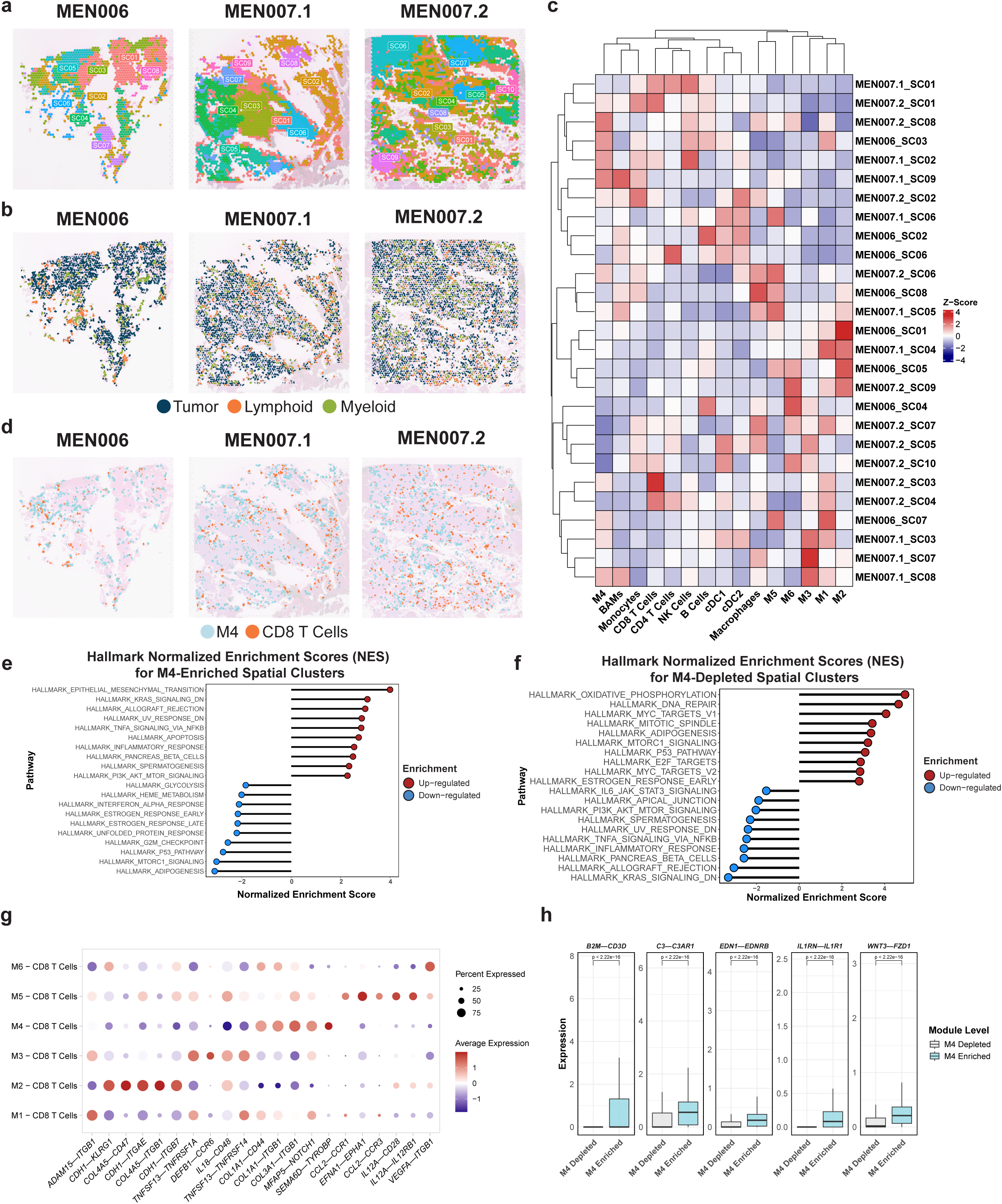
Spatial-sequencing contextualization of cell modules. **a,** Spatial transcriptomics of three meningioma samples, showing spatial cluster organization and tissue heterogeneity. **b,** CellTrek cell type mapping to spatial samples based on broad cellular categories. **c,** Heatmap of immune and tumor cell module abundance across spatial clusters of meningioma samples. **d,** CellTrek cell type mapping of EMT (M4) tumor cells and CD8 T cells in spatial samples. **e-f,** Hallmark pathway enrichment analysis, displaying normalized enrichment scores (NES) for spatial clusters stratified by M4 composition. **g,** Dot plot of ligand-receptor interactions between CD8 T cells and the 6 tumor cell modules. **h,** Box plots showing expression of ligand-receptor pairs across M4-stratified spots from spatial data, highlighting cell-cell communication specific to the M4 module.

We next sought to identify ligand-receptor interactions that might explain the observed colocalization of EMT (M4) tumor cells and CD8 T cells. Using the NICHES R package, we found that the EMT (M4) cell state primarily employed collagen protein interactions to mediate contacts with CD8 T cells (Fig. 4G). These physical contacts contrasted with the utilization of chemokine-based interactions by the other cell states, like IL12A in Stress (M5) cells, or TNFSF13 in Sterol (M3) cells. NICHES was also used to identify ligand-receptor pairs that were differentially expressed in M4-enriched spatial clusters (Fig. 4H). A subset of these, *B2M*-*CD3D, C3*-*C3AR1, EDN1*-*EDNRB, IL1RN*-*IL1R1,* and *WNT3*-*FZD1*, are highlighted in Fig. 4H, and suggest that these interactions may be correlated with the establishment of a microenvironment conducive of T cell co-localization. To draw further parallels with the L-R pairs identified at the single-cell level, the top pairs were overlaid on the spatial data and once again, there is a faithful correlation between those spatial clusters that are M4-enriched and the interactions posited to be utilized by EMT (M4) cells in their interactions with CD8 T cells (Extended Data Fig. 4E). Collectively, these data revealed that EMT (M4) tumor cells engaged in specialized ligand-receptor interactions with CD8 T cells, shaping what is likely to be a distinctive tumor microenvironment. To complement these insights, we then turned our attention to the broader immune landscape of these tumors to further elucidate how unique tumor genotypes and their corresponding phenotypes can influence immune cell function.

### Immune cells reflect unique meningioma microenvironments

To identify unique characteristics of the meningioma immune microenvironment, we compared the immune cells in our meningioma dataset with those in other intracranial tumors, the dura, and peripheral blood mononuclear cells (PBMCs). Broad categorization of immune cell subtypes revealed differences in the immune composition of these different tissue types (Fig. 5A, Extended Data Fig. 5A, B). Immune cells from intracranial tumors (Meningioma, Glioblastoma, and Vestibular Schwannoma) were primarily composed of macrophages, followed by a mix of CD8 and CD4 T cells (Fig. 5B, Supp. Table 19). The immune cells of peripheral tissues (Dura, PBMCs), on the other hand, were chiefly composed of CD4 and CD8 T cells, and the myeloid cells included both macrophages and classical monocytes.

**Fig 5.**
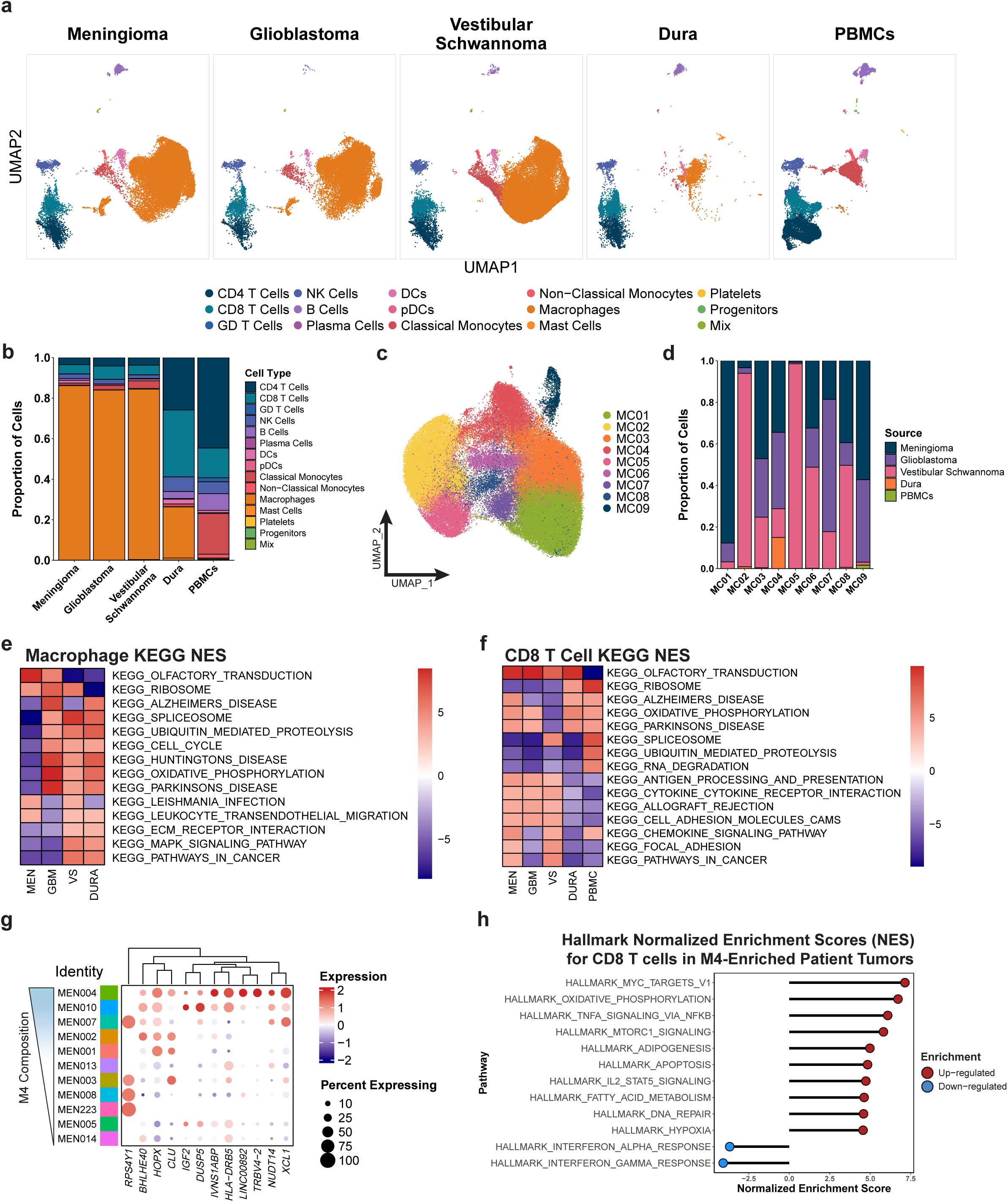
Meningioma tumors shape immune cells. **a,** UMAP visualization of immune cell populations across different histological sources, including meningioma (MEN), glioblastoma (GBM), vestibular schwannoma (VS), dura, and peripheral blood mononuclear cells (PBMCs). **b,** Bar plot showing the proportion of immune cell types across different tissue sources. **c,** De-novo UMAP clustering of macrophage cells, identifying distinct subclusters. **d,** Bar plot displaying the composition of macrophage clusters by the different histological sources. **e,** KEGG pathway enrichment analysis for macrophages across different sources. **f,** KEGG pathway enrichment analysis for CD8 T cells across different sources. **g,** Expression heatmap of identified candidate genes in CD8 T cells, ordered by increases in EMT (M4) cell state composition of tumor cells. **h,** Hallmark pathway enrichment analysis of CD8 T cells which are found in tumors that are proportionally high for the EMT (M4) cell state.

Given the frequency and predominance of macrophages, we reanalyzed these cells in isolation by performing *de-novo* normalization, harmonization, and clustering, which resulted in nine macrophage clusters, MC01-09, from all five histological sources (Fig. 5C). These nine clusters also varied in terms of their histological composition with MC02 and MC05 being primarily made up of vestibular schwannoma macrophages while meningioma macrophages made up the majority of MC01 and MC09 (Extended Data Fig. 5C). The rest of the identified macrophage clusters had variable histological compositions, with a notable case being MC04, which contained macrophages from all sources but PBMCs (Fig. 5D, Supp. Table 20). We then used gene set enrichment analysis (GSEA) to further examine large-scale, pathway-level differences between macrophages from each tissue type. We identified shared pathway utilization by macrophages from glioblastoma, vestibular schwannoma, the dura, and PBMCs. This included shared up-regulation of KEGG pathways such as ubiquitin-mediated proteolysis, cell cycle, oxidative phosphorylation, and the spliceosome (Fig. 5E). However, meningioma macrophages were functionally distinct from the others as illustrated by the downregulation of these pathways and select up-regulation of olfactory transduction and ribosome KEGG pathways. Unique pathway utilization by meningioma macrophages was observed across multiple gene sets (Extended Data Fig. 5D-E, Supp. Table 21) and further underscored the distinct immune microenvironment of this tumor type.

We also carried out gene set enrichment analysis for CD8 T cells across histological sources, because this population of cells exhibited a pronounced association with the microenvironmentally-modulating M4 cell state (Fig. 5F, Supp. Table 22). This again revealed a pathway utilization signature unique to meningioma, but to a lesser degree than macrophages, with intratumoral CD8 T cells utilizing pathways including olfactory transduction, cytokine to cytokine receptor interaction, and antigen processing and presentation. This contrasted starkly with CD8 T cell pathway utilization in PBMCs, which reflected an active, circulating phenotype via upregulation of KEGG pathways relating to the ribosome, spliceosome, ubiquitin-mediated proteolysis, and RNA degradation. Interestingly, the CD8 T cells of the dura exhibited an intermediate pathway utilization phenotype, with overlap in up- and down-regulated pathways from both intratumoral and PBMC CD8 T cells.

To expand on the spatial and compositional correlations between CD8 T cells and the EMT (M4) cell state, we stratified our patients based on the M4 composition of their tumors. In doing so, we identified two patients, MEN004 and MEN010, as M4-high, with tumors composed of 92.45% and 59.57% M4 cells, respectively (Extended Data Fig. 5F). The rest of the patients were classified as M4-low, with M4 proportions not exceeding 27.38% of the malignant population. Differential gene expression analysis was performed on T-cells from these two groups, and the genes defining T-cells from M4-high and M4-low tumors were visualized as a volcano plot (Extended Data Fig. 5G, Supp. Table 23). A subset of these genes was then used to generate a clustered heatmap to better illustrate these relationships (Fig. 5G). Among the genes upregulated with increases in M4 composition were *BHLHE40* and *HOPX,* which are factors known to be involved in not only CD8 T cell differentiation but also exhaustion. We also see increases in expression of *XCL1*, highlighting the functional recruitment of other immune cells^36^ by these T cells. To contextualize individual genes, we then performed gene set enrichment analysis on CD8 T cells stratified by tumoral M4 composition (Fig. 5H, Supp. Table 24). CD8 T cells from M4-high tumors up-regulate pathways indicative of a stressful microenvironment such as apoptosis, hypoxia, and *TNFa* signaling via *NFKB*. These cells also show concomitant down-regulation of interferon alpha and gamma responses, likely indicative of a dysfunctional phenotype. Taken together, our findings demonstrate that the meningioma microenvironment fosters distinctive immune cell phenotypes, particularly among macrophages and CD8 T cells, the latter of which illustrated the interplay between our tumor genotype-driven cell states and the immune milieu.

## Discussion

Recent studies have shown that meningiomas can be classified based on shared genetic and epigenetic alterations, or by methylation-based classifiers^16,37,38^. Compared to the WHO histopathological subtypes, these molecular classifiers better predict clinical behavior, including risk of recurrence and overall patient outcomes^3,39^. However, these classifications do not neatly map one-to-one across publications or molecular modalities, highlighting the complex heterogeneity within and between meningiomas, and the nuanced relationships among genomic data types. Here, we use multi-modal analysis—combining scRNA-seq, exome sequencing, bulk RNA-seq, and spatial transcriptomics—to uncover novel relationships between transcriptional heterogeneity in meningiomas and their underlying genotypes, chromosomal alterations, immune microenvironments and potential cell of origin.

In our cohort, we identified six overarching tumor cell states, M1-M6, that represent distinct transcriptional programs and exhibit associations with previously recognized genetic drivers. For instance, the Sterol Metabolism (M3) module was linked to *TRAF7, KLF4* and *AKT1* mutations, whereas the EMT (M4) module was associated with *NF2* mutations and Chr22q loss. These findings align with the established large-scale genomic distinction between *NF2*-driven and non- *NF2* meningiomas^9,33^. By utilizing scRNA-seq as our primary experimental tool, we characterize heterogeneity that might otherwise be obscured in other analyses given that even tumors within the same genetic class can harbor multiple tumor cell populations^40^.

We also observed direct relationships between tumor cell states and the immune microenvironment. Bulk RNA-sequencing revealed specific patterns of immune infiltration associated with the EMT (M4) tumor cell state, which were corroborated by spatial transcriptomics data. However, these immune cells displayed characteristics of dysfunction, mirroring findings in other collagen-rich or fibrotic tumors, where dense extracellular matrices can hinder immune cell activity^41–43^. In meningiomas, such collagen-driven niches associated with our M4 cell state may be particularly relevant given the frequent co-occurrence of the fibrous histology with Chr22q loss and *NF2* inactivation^15^.

Recent large-scale genomic studies have also described transcriptional and epigenetic programs underlying meningioma biology. The six tumor cell states we identified partially align with these published frameworks, with two in particular, M3 and M4, most closely mirroring published transcriptional programs. The sterol metabolism state (M3) is highly enriched for *TRAF7*/*KLF4*/*AKT1* mutations, oxidative phosphorylation pathways, and lipid handling genes which resemble the *NF2*-wildtype MG2 subgroup from Nassiri et al.^3^, the Type A tumors of Patel et al^13^., and the “Merlin-intact” methylation class of Choudhury et al^16^. Like those groups, the M3 tumor cell state we identified tends to associate with lower histological grades and lower recurrence rates. However, the methylation-defined angiogenic signature is not expressed as strongly in our M3 cell state, in favor of a metabolic profile centered around fatty acid and sterol metabolism.

The EMT cell state (M4), characterized by *NF2*/Chr22q loss, displays marked collagen remodeling and a strongly immune-enriched profile. This parallels the “immunogenic” MG1 group, the Type B tumors, and the “Immune-enriched” methylation class. All of these groups show convergent genomic instability with frequent co-occurrence of *NF2* mutations and Chr22q loss with overarching regulation of immune signaling pathways and enrichment of immune cells. The association with histological grade and recurrence is discordant amongst these groups, with some displaying better prognosis (MG1, Nassiri), and others, including our M4 cell state, displaying a more nuanced and intermediate prognosis (Immune-enriched, Choudhury). The other cell states in our data (M1, M2, M5, M6) do not have marked associations with mutations or chromosomal aberrations that fit into the canonically established subgroups. For example, the M1 cell state is associated with both losses in Chr1p and mutations in *KLF4*. However, phenotypically, this cell state shows upregulation of cell cycling and mitotic genes, most similar to that found in the Proliferative MG4 subgroup, Type C tumors, and the Hypermitotic methylation class, rather than one associated with *KLF4* mutations. These more nuanced cell states are further exemplified by the M2, M5 and M6 cell states, which correspond to chromatin remodeling, stress responses and hypoxia, respectively. These states likely represent more generalized cellular processes that are agnostic to tumor genetics, but still contribute to shaping tumor cell phenotypes.

Our results suggest that meningioma tumor cell phenotypes are not strictly defined by genetics, and point to potential differences in cell of origin. By mapping individual cells to curated transcriptional signatures for each meningeal layer (dura, pia, arachnoid), we observed that individual tumors tended to resemble one layer more strongly than the others. This suggests that meningiomas may originate from region-restricted progenitors that retain some “memory” of that layer even after malignant transformation. Conversely, we also observe that genetically similar tumors, such as those that were NF2-driven, can transcriptionally resemble different layers, implying that driver mutations intersect with cell-of-origin to shape downstream transcriptional programs and immune niches. This suggests that meningioma heterogeneity may stem not only from genotype but also from anatomically distinct progenitors.

Overall, our results corroborate the current understanding that meningioma biology is largely governed by distinct genetic drivers, and demonstrate that these drivers impart unique tumor cell states and immune microenvironments. By characterizing distinct tumor cell states at single-cell resolution, we provide evidence that *NF2* and non-*NF2* driven tumors give rise to different transcriptional and immune landscapes. These findings complement existing genomic and methylation-based classifiers as we continue to learn more about how tumor genotypes and their corresponding cell states influence tumor behavior and progression. Future work utilizing larger patient cohorts and additional modalities (e.g. methylation) will be required to determine whether these cell states correlate with clinical outcomes or treatment responses, and how they might be integrated into both established and novel classification frameworks.

## Methods

### Patients and samples

Adult patients undergoing neurosurgical intervention at Barnes-Jewish and Massachusetts General Hospital were screened. Selection criteria included (1) age > 18 years and (2) presence of intracranial meningioma with clinical indications for surgical resection. All samples were collected from patients undergoing surgical resection for a primary meningioma tumor. Prior to surgery, informed consent was obtained from patients meeting selection criteria following the Washington University School of Medicine Institutional Review Board Protocol #202107071, the Massachusetts General Hospital IRB Protocol #2022P001982. During surgical resection, specimens were placed in normal saline and immediately maintained on ice pending further processing. Clinical characteristics are summarized in Supp. Table 1. All procedures and experiments were performed in accordance with the Helsinki Declaration.

### Next-generation sequencing of meningioma DNA samples

Total RNA and genomic DNA was co-purified from fresh-frozen tumor tissue using the AllPrep Mini RNA/DNA kit (catalog #80204, Qiagen, Valencia, CA). Manufacturer’s instructions were modified to incorporate a 5 minute incubation for the RNA and DNA elution. The tissue was homogenized and lysed using the Tissue Lyser II (Qiagen) set to 45 Hrz for 2 minutes. Additionally, DNA was purified from frozen blood using the QIA-AMP Blood and Tissue kit according to manufacturer’s instructions (catalog # 51104, Qiagen, Valencia, CA). (Quantity assessment was determined using the Qubit and corresponding quantification kits. (Invitrogen, Carlsbad, CA)

### Variant detection in DNA sequencing

Exome sequencing data for each tumor-normal (blood) pair from each patient was aligned to hg38 reference genome using the “Burrows–Wheeler” aligner with the maximal exact matches (BWA-MEM) algorithm^44^. Aligned reads were then sorted, deduplicated, and processed through base quality score recalibration (GATK)^45^. Structural variants and large insertions or deletions were identified using Manta^46^, while single nucleotide variants (SNVs) and small indels were detected using VarScan2^47^, Strelka2^48^, and MuTect2^49^.SNV/indel annotation was performed using Variant Effect Predictor (VEP), version 95^50^. Variants with a population frequency >0.1% in gnomAD were excluded, as well as those located in regions with low mapping quality or insufficient coverage (<20×). The complete somatic variant analysis pipeline is publicly available as a Common Workflow Language (CWL) workflow at https://github.com/genome/analysis-workflows (Commit ID: 03650ca89aca9b4edfeb213f720f154f0ed18f10; https://github.com/genome/analysis-workflows/commit/03650ca89aca9b4edfeb213f720f154f0ed18f10). Copy number variants (CNVs) were inferred and visualized using CNVkit^51^.

### Analysis of scRNA-seq 10X data

To generate single-cell suspensions of freshly resected primary meningioma samples were placed on ice and macerated into a paste and enzymatically digested using a human tumor dissociation kit (Miltenyi Biotec). Following dissociation, the samples were resuspended in 5 mL of trypsin for 3-4 minutes to further dissociate the collagen-rich tumors. After this secondary dissociation step, the tumor digest was resuspended in FBS-supplemented media and pelleted. Afterwards, the sample was resuspended in ACK lysis buffer (Lonza) for lysis of remaining red blood cells. The resulting cell suspension was evaluated under a microscope for the significant presence of debris. If the suspension had considerable debris, samples were subjected to a debris removal solution according to the manufacturer’s instructions (Miltenyi Biotec). Cells were resuspended in 10% FBS/IMDM in preparation for construction of single-cell libraries according to the manufacturer’s protocol (10x Genomics) for 5’ sequencing. The established libraries were sequenced on a NovaSeq6000 S4 Flow Cell using the XP workflow with a sequencing depth of 50,000 reads per cell and a 151×10×10×151 sequencing recipe according to manufacturer protocol for 5′ sequencing. The resulting FASTQ files were then processed using the CellRanger analysis suite (https://github.com/10XGenomics/cellranger) with alignment to hg38 reference genome. Cells were filtered with a minimum feature threshold of 1000, a maximum count threshold of 93 percentile, and mitochondrial percentage of less than 5%. We used multiple doublet removal algorithms, and removed a cell if it was called positive of any of the algorithms. We then ran samples through CellBender (https://github.com/broadinstitute/CellBender) to minimize technical artifacts, specifically ambient RNA and random barcode swapping from UMI-based count matrices^52^. Using the Seurat package v4.0.3, data was log normalized, and the top 2000 variable features were selected and scaled with regression of UMI counts^53,54^. Principal component analysis and uniform manifold approximation project were run using 30 principal components. A K-nearest neighbors graph was generated using default settings of the Seurat function FindNeighbors and Louvain clustering was implemented using FindClusters with a resolution of 0.6. Small (<100 cells) Louvain-based clusters distinct from neoplastic cells and expressing macrophage markers were removed. Integration was performed by splitting the dataset by patient, normalizing, scaling and regressing by UMI count, running PCA, then using FindIntegrationAnchors and IntegrateData for Reciprocal PCA (RPCA).

### CNV identification with CONICS

Along with CNV identification from WES, we used CONICS to identify CNVs in our 10X scRNA-seq data as outlined at: (https://github.com/diazlab/CONICS)^55^. For further characterization of CNVs and their association with tumor cell states, the posterior probability score was utilized to assign each cell to a particular CNV. Hard calls were produced by binarizeMatrix() using the default thresholds (probability > 0.8 for gains, < 0.2 for losses).

### Identification of differentially expressed genes

Differentially expressed genes were found using the default settings of the wilcoxauc function from the Presto R package (https://github.com/immunogenomics/presto/). Top DEGs for CNV analysis, NMF modules, gene set enrichment analysis, and immune cell types were found by filtering for genes with p_adj ≤ 0.05 and sorting by a combination of highest LogFC value, lowest adjusted p-value and largest percent expression differences between groups.

### Non-negative matrix factorization (NMF) of scRNA-seq count matrix

We performed NMF on each tumor to capture intratumoral heterogeneity within patients. Negative values in the scaled matrix were set to zero, and K values ranging from 4-9 were used on each patient as in Gavish et al.^56^. To find robust programs, multiple filtering parameters were iteratively applied to the factorized matrix programs using programs of various sizes. A minimum overlap (intra_min) with a program from the same cell line was applied to select for robustness. A maximum overlap (intra_max) with a program from the same cell line was applied to remove redundant programs. A minimum overlap (inter_min) with a program from another cell line was also applied to reduce sparsity and enhance modularity in the programs. We found that an intra_min=35, intra_max=5, and inter_min=12 using programs composed of 75 genes created high complexity correlation and robust gene programs that included all patients or most of the tumors. Programs with high correlation to each other and that were shared across multiple patients were combined into modules. Six modules were identified, and gene sets composed of the combined programs were filtered by taking genes that had an NMF score of higher than 0.25. Functional enrichment was performed on the gene sets from each module using Topfunn and CytoScape (https://cytoscape.org/). The addModuleScore() function^57^ was used to score each cell with an average expression of that module. Cells were then categorized into a module by which module had the maximum score. Downstream utilization of these modules scores include comparison across the derived CNVs as identified by CONICs or functional enrichment stratified by fGSEA.

### Gene set enrichment analysis (GSEA)

For large-scale pathway utilization analyses, we used the fGSEA R package to test for enrichment of multiple genesets downloaded from MsigDB including Hallmark, KEGG, GO:BP, GO:CC, GO:MF, and WikiPathways^58^. For input, we used either *z*-score statistics from Seurat DEG analysis or pre-ranked gene lists generated using a fast Wilcoxon rank-sum test from the presto R package.

### Potency determination using CytoTRACE2

To characterize the “developmental potency” of the meningioma tumor cells, we applied the scoring algorithm from the CytoTRACE2 package to our unintegrated tumor cell Seurat object by executing the cytotrace2 function using default parameters^34^.

### Bulk RNA-seq deconvolution and correlation analysis from Patel et al

Bulk RNA-sequencing of primary sporadic meningiomas was obtained from Gene Expression Omnibus (GEO) accession GSE136661^13^ and genes were annotated using reference version Human.GRCh38.p13. To quantify the cell fractions of our NMF-derived gene signatures (M1-M6) and the diverse set of immune infiltrates in the bulk samples, we used the online bulk deconvolution platform CIBERSORTx^59^. For this analysis, all primary meningioma samples were used regardless of mutational status, since we wanted our analysis to be applicable to the diversity of genomic variation seen in the clinic. Normalized counts of our meningioma sc-RNAseq dataset were used to generate the signature matrix file, while normalized bulk counts were used as the mixture file. CIBERSORTx was then run using default parameters and the resulting fraction matrix was used for downstream analyses. Correlations between NMF module composition for patient tumors and between clinical variables (WHO grade, sex, recurrence, age) were assessed. Pearson correlation coefficients and p-values were calculated (Supp. Table 25). For continuous clinical variables (age), an R-squared value was also calculated.

### Spatial transcriptomic analysis

Using the Seurat package’s Load10x_Spatial function, filtering parameters of nCount_Spatial > 500 & nFeature_Spatial > 200, and SCTransform normalization, we derived spatial transcriptomic samples that ranged from 1281 to 3414 spots. After running PCA, clusters were identified using 1:30 dimensions and spatially variable features were found via the FindSpatiallyVariableFeatures function. The resulting object served as the basis for downstream analysis using the CellTrek package^60^. To co-embed the spatial and our meningioma single-cell transcriptomic dataset, we first used the traint function with default parameters. Following this, the single cells from our established reference were then mapped to their spatial location via the celltrek function using the following parameters: intp=T, intp_pnt=5000, intp_lin=F, nPCs=30, ntree=1000, dist_thresh=0.55, top_spot=10, spot_n=5, repel_r=20, repel_iter=20, keep_model=T. The resulting object was used for downstream analyses. For spatial cluster composition analysis, each cell that was co-embedded and charted to a spatial location was then mapped back on to the spot it was closest to using the FNN package’s get.knnx function^61^. Once mapped back to the spatial spots, these cells were then enumerated and clusters were tabulated for composition across all three samples and compared via z-scoring for each cell type.

### Cross-platform ligand-receptor analysis using NICHES

We applied the established ligand-receptor signaling determination pipeline established in the NICHES package on both our scRNA-seq and spatial transcriptomic data to generate an effective cell-cell signaling repertoire for meningioma cells^62^. In both instances, the runNICHES function was run with an imputed version of either the single-cell or spatial Seurat objects to amplify low-strength signals and remove gene-sampling-related clustering artifacts. In all cases, the assay used was the “alra” assay, and the LR.database was “fantom5”. The resulting NICHES output was then either visualized as a heatmap or dotplot by extracting the data from the “ReceivingType” metadata. To identify the most differentially expressed ligand-receptor pairs, the FindAllMarkers function was used with “roc” as the test being used and top pairs were selected by filtering based on decreasing “myAUC” values.

### Statistical Analysis

For bulk sequencing comparisons of continuous groups, Spearman correlations were used. For comparisons between continuous and categorical variables, a point biserial correlation was used. For comparisons between two categorical variables, a phi coefficient was calculated.

Adjusted p-values were derived using a Benjamini-Hochberg procedure. For comparison of groups, For the comparison of groups, the Mann–Whitney U test was used. For differential gene expression, the Wilcoxon rank-sum test (two-sided) was used with Bonferroni correction. P ≤ 0.05 was considered to be statistically significant. Statistical analyses were performed with R statistical language V 4.1.0 or Seurat.

### Data Availability

FASTQ files will be available in the NCBI Sequence Read Archive (SRA) and processed gene expression and Seurat objects used for all analyses will be available on the open-access data-sharing platform Zenodo (RRID: SCR_004129) upon publication.

---

**Extended Data Fig 1.**
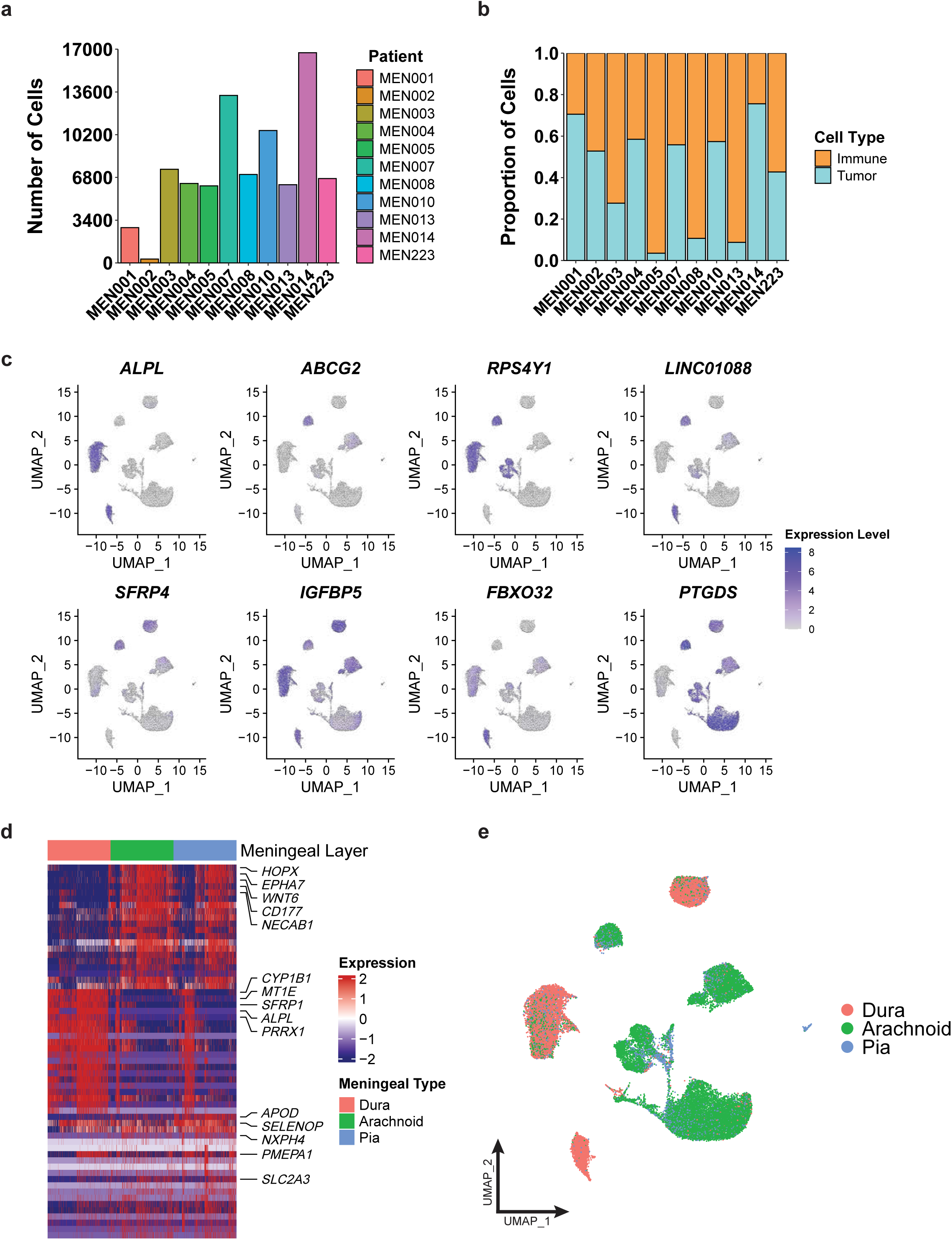
Patient heterogeneity alludes to cell-of-origin differences. **a,** Bar plot showing the total number of cells (both tumor and immune) from individual patients. **b,** Bar plot showing the proportion of cells assigned to each cellular category (tumor, immune) across meningioma tumors from individual patients. **c,** Feature plots displaying expression level of patient-specific genes across all tumor cells. **d,** Heatmap of differentially expressed genes for cells assigned to each meningeal layer; Hierarchical clustering of gene expression across meningeal layers, with expression levels scaled from low (blue) to high (red). **e,** UMAP visualization of single-cell transcriptomes, with cells colored by assigned meningeal layer.

**Extended Data Fig 2.**
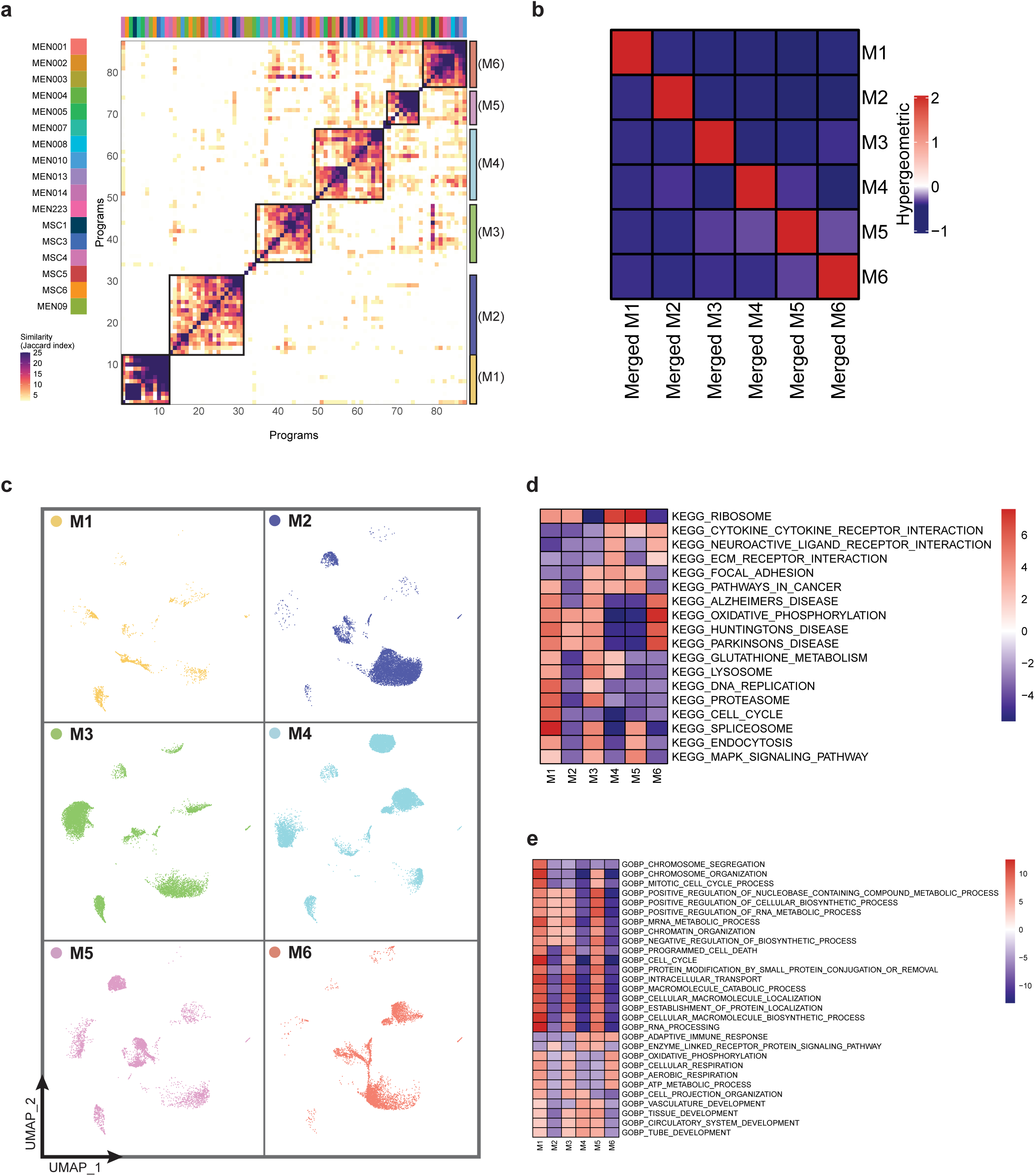
Tumor cell modules provide a robust framework for biological differences. **a,** Clustered heatmap of transcriptional programs derived from NMF across meningioma samples including external datasets, highlighting the robust nature of the modules across meningiomas. **b,** Hypergeometric heatmap illustrating the overlap between the cell module (M1-M6) from our initial NMF determination and those found from the merged datasets. **c,** UMAP visualization of single-cell transcriptomes, with cells colored and split by assigned transcriptional modules. **d-e,** KEGG and GO:BP pathway enrichment analysis for tumor cells across different cell states.

**Extended Data Fig 3.**
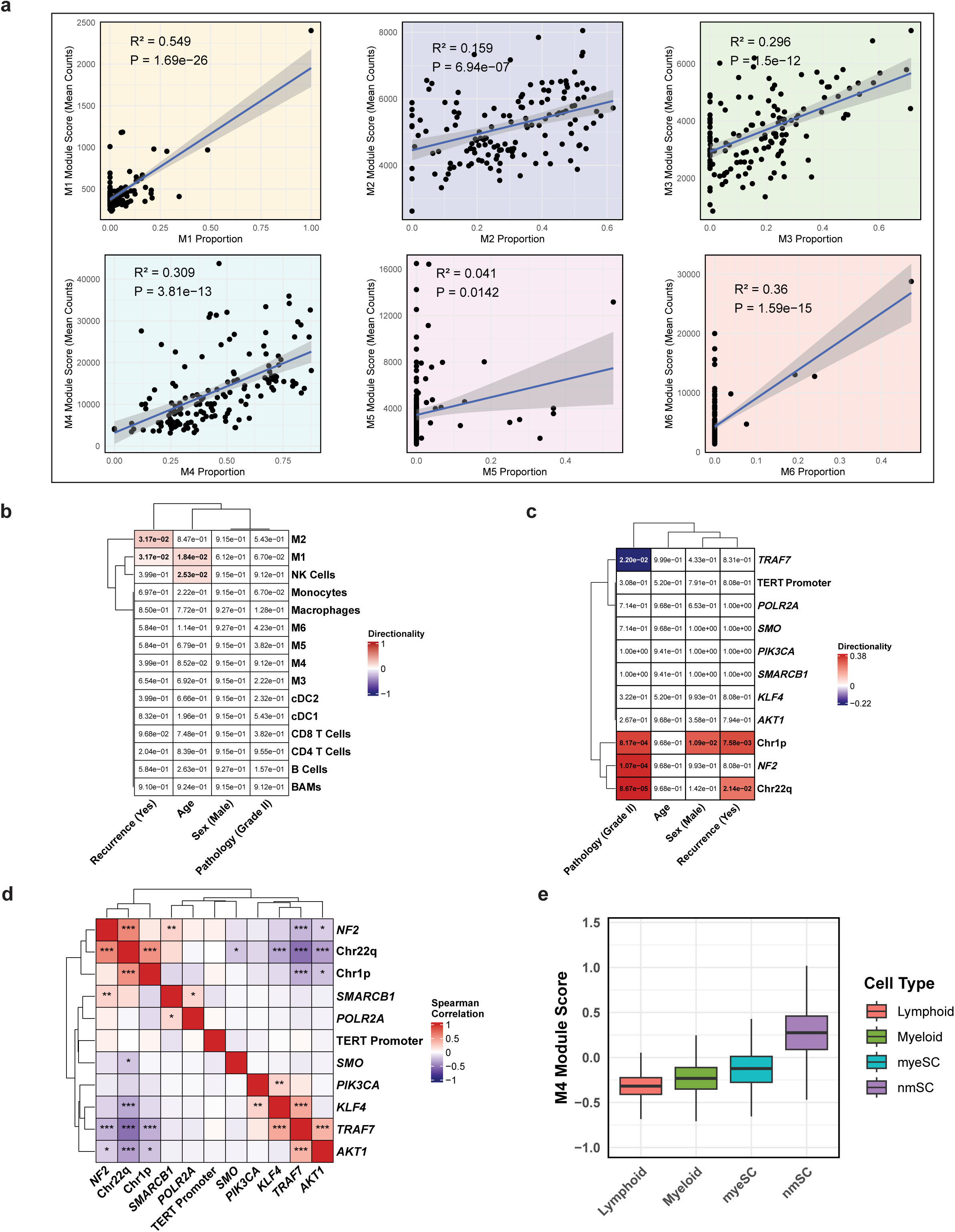
Bulk-RNA correlations expand dichotomy between Sterol (M3) and EMT (M4) tumor cell states. **a,** Module score correlation with cell state composition in bulk RNA-seq samples. **b-c,** Correlations between cell type (b) or mutations (c) with the corresponding clinical variables in bulk RNA-seq data, with directionality scaled from negative (blue) to positive (red) and p-values displayed. **d,** Correlations amongst mutations from Patel data, with Spearman correlation values scaled from -1 (blue) to +1 (red) and p-values summarized (*** = p-value < 0.001, ** = p-value < 0.01, * = p-value < 0.05). **e,** Box plots displaying EMT (M4) module scores for distinct cell types in vestibular schwannoma tumors (myeSC = myelinating Schwann cells, nmSC = non-myelinating Schwann cells).

**Extended Data Fig 4.**
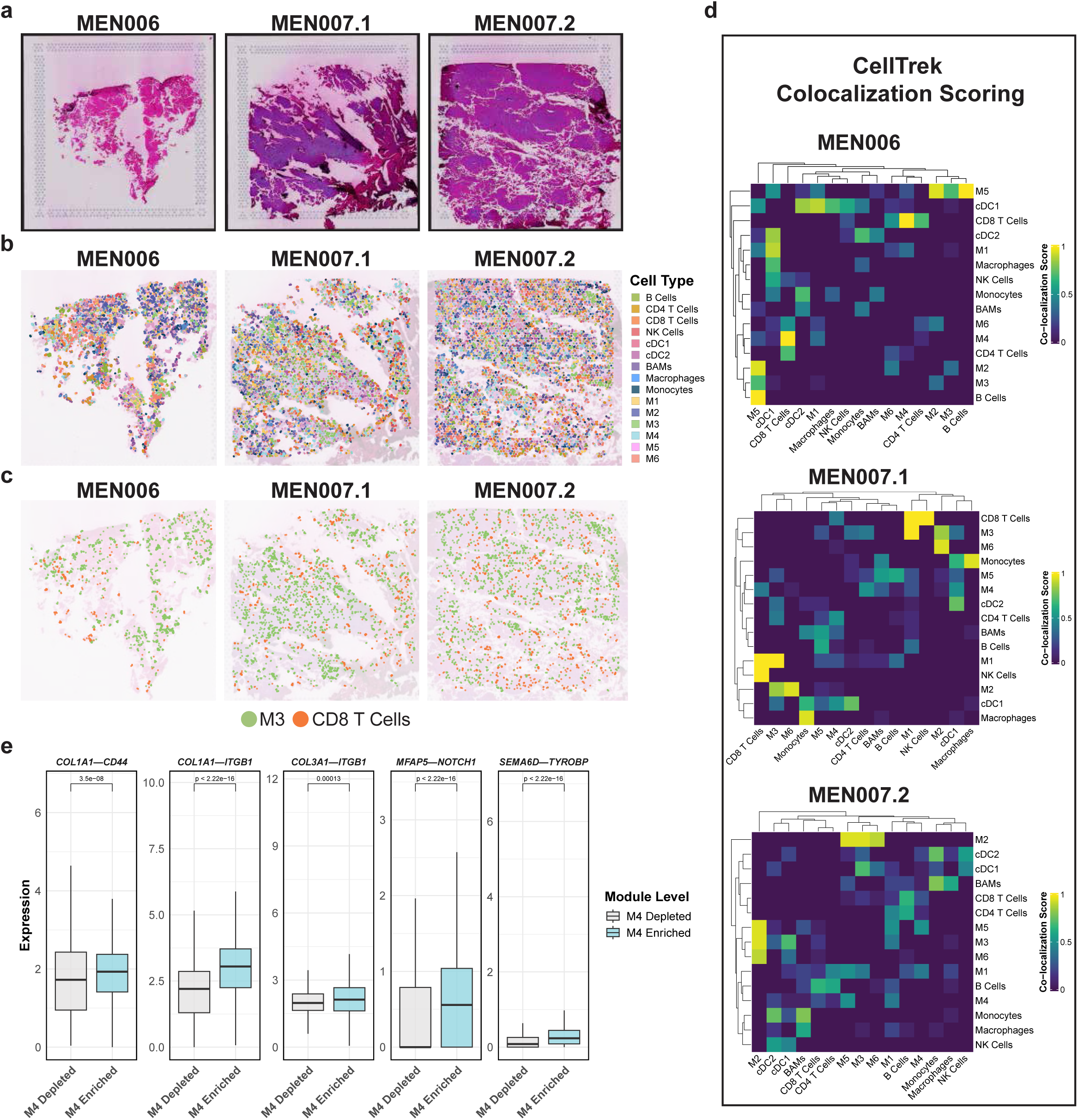
Spatial transcriptomics corroborate co-localization patterns. **a,** Hematoxylin and Eosin (H&E) staining of the three spatial samples, MEN006, MEN007.1 and MEN007.2. **b,** CellTrek mapping of specific cell types overlaid on H&E images of spatial samples. **c,** CellTrek mapping of Sterol (M3) tumor cells and CD8 T cells overlaid on H&E images of spatial samples. **d,** Individual co-localization score heatmaps derived by CellTrek. **e,** Box plots showing expression of ligand-receptor pairs derived from single-cell NICHES analysis across M4-stratified spots from spatial data, highlighting multi-platform convergence.

**Extended Data Fig 5.**
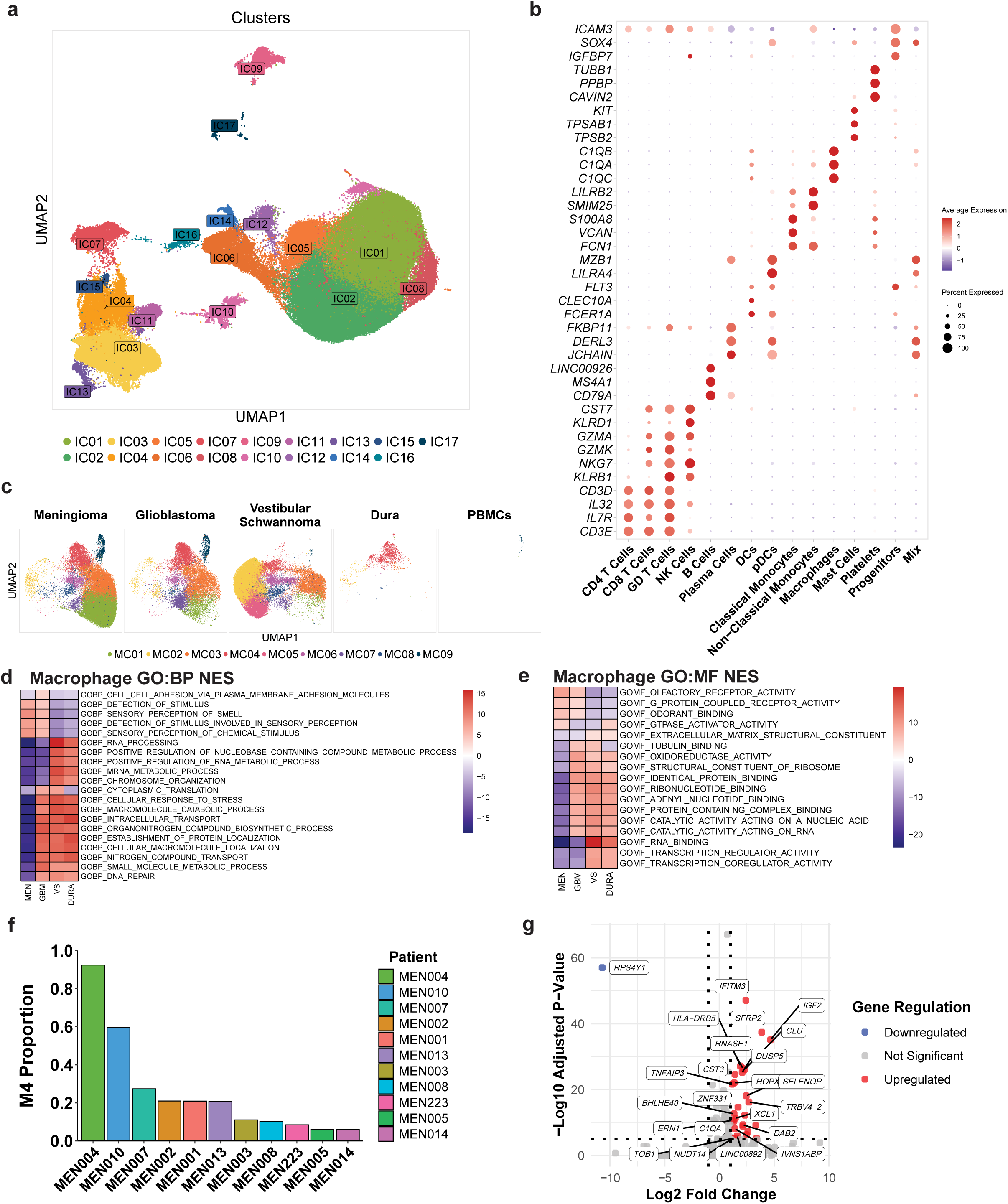
Histology-specific immune modulation. **a,** UMAP visualization of immune cell transcriptomes from different histological sources, with cells colored by cluster. **b,** Dot plot displaying immune cell-specific gene expression patterns across all histological sources. **c,** UMAP visualization of macrophage single-cell transcriptomes, with cells colored by macrophage cluster and split by histological source. **d-e,** GO:BP and GO:MF pathway enrichment analysis for macrophages across different histological sources. **f,** Bar plot showing the proportion of EMT (M4) tumor cells in individual patients’ meningioma tumors. **g,** Volcano plot illustrating the genes that were selectively up- and down-regulated based on M4-enrichment (p-value < 0.05, Log2 Fold Change > 1).

## References

1. Ostrom, Q. T., et al. CBTRUS Statistical Report: Primary Brain and Other Central Nervous System Tumors Diagnosed in the United States in 2012-2016. Neuro. Oncol. 21, v1–v100 (2019).

2. Maas, S. L. N. et al. Integrated Molecular-Morphologic Meningioma Classification: A Multicenter Retrospective Analysis, Retrospectively and Prospectively Validated. J. Clin. Oncol. 39, 3839–3852 (2021).

3. Nassiri, F. et al. A clinically applicable integrative molecular classification of meningiomas. Nature 597, 119–125 (2021).

4. Driver, J. et al. A molecularly integrated grade for meningioma. Neuro. Oncol. 24, 796–808 (2022).

5. Patel, B. et al. Identification and management of aggressive meningiomas. Front. Oncol. 12, 851758 (2022).

6. Perry, A., Scheithauer, B. W., Lohse, C. M. & Wollan, P. C. ‘Malignancy’ in meningiomas. Cancer 85, 2046–2056 (1999).

7. Okano, A. et al. Associations of pathological diagnosis and genetic abnormalities in meningiomas with the embryological origins of the meninges. Sci. Rep. 11, 6987 (2021).

8. Brastianos, P. K. et al. Genomic sequencing of meningiomas identifies oncogenic SMO and AKT1 mutations. Nat. Genet. 45, 285–289 (2013).

9. Clark, V. E. et al. Genomic analysis of non-NF2 meningiomas reveals mutations in TRAF7, KLF4, AKT1, and SMO. Science 339, 1077–1080 (2013).

10. Choy, W. et al. The molecular genetics and tumor pathogenesis of meningiomas and the future directions of meningioma treatments. Neurosurg. Focus 30, E6 (2011).

11. Reuss, D. E. et al. Secretory meningiomas are defined by combined KLF4 K409Q and TRAF7 mutations. Acta Neuropathol. 125, 351–358 (2013).

12. Youngblood, M. W. et al. Correlations between genomic subgroup and clinical features in a cohort of more than 3000 meningiomas. J. Neurosurg. 133, 1345–1354 (2020).

13. Patel, A. J. et al. Molecular profiling predicts meningioma recurrence and reveals loss of DREAM complex repression in aggressive tumors. Proc. Natl. Acad. Sci. U. S. A. 116, 21715–21726 (2019).

14. Louis, D. N. et al. The 2021 WHO classification of tumors of the Central Nervous System: A summary. Neuro. Oncol. 23, 1231–1251 (2021).

15. Sahm, F. et al. DNA methylation-based classification and grading system for meningioma: a multicentre, retrospective analysis. Lancet Oncol. 18, 682–694 (2017).

16. Choudhury, A. et al. Meningioma DNA methylation groups identify biological drivers and therapeutic vulnerabilities. Nat. Genet. 54, 649–659 (2022).

17. Nassiri, F. et al. DNA methylation profiling to predict recurrence risk in meningioma: development and validation of a nomogram to optimize clinical management. Neuro. Oncol. 21, 901–910 (2019).

18. Lucas, C.-H. G. et al. Spatial genomic, biochemical and cellular mechanisms underlying meningioma heterogeneity and evolution. Nat. Genet. 56, 1121–1133 (2024).

19. Fan, H. et al. Decoding meningioma heterogeneity and neoplastic cell-macrophage interaction through single-cell transcriptome profiling across pathological grades. J. Transl. Med. 21, 751 (2023).

20. Vasudevan, H. N. et al. Intratumor and informatic heterogeneity influence meningioma molecular classification. Acta Neuropathol. 144, 579–583 (2022).

21. Bayley, J. C., 5th et al. Multiple approaches converge on three biological subtypes of meningioma and extract new insights from published studies. Sci Adv 8, eabm6247 (2022).

22. Xu, D., Yin, S. & Shu, Y. NF2: An underestimated player in cancer metabolic reprogramming and tumor immunity. NPJ Precis. Oncol. 8, 133 (2024).

23. Lin, J. et al. IGFBP5, as a Prognostic Indicator Promotes Tumor Progression and Correlates with Immune Microenvironment in Glioma. J. Cancer 15, 232–250 (2024).

24. Chang, X. et al. Genomic and transcriptome analysis revealing an oncogenic functional module in meningiomas. Neurosurg. Focus 35, E3 (2013).

25. Kalamarides, M. et al. Identification of a progenitor cell of origin capable of generating diverse meningioma histological subtypes. Oncogene 30, 2333–2344 (2011).

26. Yamashima, T. et al. Prostaglandin D synthase (beta-trace) in human arachnoid and meningioma cells: roles as a cell marker or in cerebrospinal fluid absorption, tumorigenesis, and calcification process. J. Neurosci. 17, 2376–2382 (1997).

27. Degenhard, T. et al. An integrative layer-resolved atlas of the adult human meninges. bioRxiv 2024.11.13.623440 (2024) doi:10.1101/2024.11.13.623440.

28. Wang, A. Z. et al. Single-cell profiling of human dura and meningioma reveals cellular meningeal landscape and insights into meningioma immune response. Genome Med. 14, 49 (2022).

29. Wolffe, A. P. Chromatin remodeling: why it is important in cancer. Oncogene 20, 2988–2990 (2001).

30. Lu, N. et al. CSMD3 is associated with tumor mutation burden and immune infiltration in ovarian cancer patients. Int. J. Gen. Med. 14, 7647–7657 (2021).

31. Swierczynski, J., Hebanowska, A. & Sledzinski, T. Role of abnormal lipid metabolism in development, progression, diagnosis and therapy of pancreatic cancer. World J. Gastroenterol. 20, 2279–2303 (2014).

32. Aouizerat, B. E. et al. A genome scan for familial combined hyperlipidemia reveals evidence of linkage with a locus on chromosome 11. Am. J. Hum. Genet. 65, 397–412 (1999).

33. Nouri, S. H. et al. Role of NF2 mutation in the development of eleven different cancers. Cancers (Basel) 17, 64 (2024).

34. Kang, M. et al. Mapping single-cell developmental potential in health and disease with interpretable deep learning. bioRxivorg 2024.03.19.585637 (2024) doi:10.1101/2024.03.19.585637.

35. Barrett, T. F. et al. Single-cell multi-omic analysis of the vestibular schwannoma ecosystem uncovers a nerve injury-like state. Nat. Commun. 15, 478 (2024).

36. Kamei, M. et al. Intratumoral delivery of a highly active form of XCL1 enhances antitumor CTL responses through recruitment of CXCL9-expressing conventional type-1 dendritic cells. Int. J. Cancer 154, 2176–2188 (2024).

37. Olar, A. et al. Global epigenetic profiling identifies methylation subgroups associated with recurrence-free survival in meningioma. Acta Neuropathol. 133, 431–444 (2017).

38. Robert, S. M. et al. The integrated multiomic diagnosis of sporadic meningiomas: a review of its clinical implications. J. Neurooncol. 156, 205–214 (2022).

39. Lee, Y. et al. Genomic landscape of meningiomas. Brain Pathol. 20, 751–762 (2010).

40. Wu, J. K. et al. Clonal analysis of meningiomas. Neurosurgery 38, 1196–200; discussion 1200–1 (1996).

41. Rømer, A. M. A., Thorseth, M.-L. & Madsen, D. H. Immune modulatory properties of collagen in cancer. Front. Immunol. 12, 791453 (2021).

42. Peng, D. H. et al. Collagen promotes anti-PD-1/PD-L1 resistance in cancer through LAIR1-dependent CD8+ T cell exhaustion. Nat. Commun. 11, 4520 (2020).

43. Larsen, A. M. H. et al. Collagen density modulates the immunosuppressive functions of macrophages. J. Immunol. 205, 1461–1472 (2020).

44. Li, H. Aligning sequence reads, clone sequences and assembly contigs with BWA-MEM. arXiv [q-bio.GN] (2013).

45. van der Auwera, G. & O’Connor, B. D. Genomics in the Cloud: Using Docker, GATK, and WDL in Terra. (O’Reilly Media, Incorporated, 2020).

46. Chen, X., et al. Manta: rapid detection of structural variants and indels for germline and cancer sequencing applications. Bioinformatics 32, 1220–1222 (2016).

47. Koboldt, D. C. et al. VarScan 2: somatic mutation and copy number alteration discovery in cancer by exome sequencing. Genome Res. 22, 568–576 (2012).

48. Kim, S. et al. Strelka2: fast and accurate calling of germline and somatic variants. Nat. Methods 15, 591–594 (2018).

49. Cibulskis, K. et al. Sensitive detection of somatic point mutations in impure and heterogeneous cancer samples. Nat. Biotechnol. 31, 213–219 (2013).

50. McLaren, W. et al. The Ensembl Variant Effect Predictor. Genome Biol. 17, 122 (2016).

51. Talevich, E., Shain, A. H., Botton, T. & Bastian, B. C. CNVkit: Genome-Wide Copy Number Detection and Visualization from Targeted DNA Sequencing. PLoS Comput. Biol. 12, e1004873 (2016).

52. Fleming, S. J. et al. Unsupervised removal of systematic background noise from droplet-based single-cell experiments using CellBender. Nat. Methods 20, 1323–1335 (2023).

53. Hao, Y. et al. Integrated analysis of multimodal single-cell data. Cell 184, 3573–3587.e29 (2021).

54. Hao, Y. et al. Dictionary learning for integrative, multimodal and scalable single-cell analysis. Nature Biotechnology 42, 293–304 (2023).

55. Müller, S., Cho, A., Liu, S. J., Lim, D. A. & Diaz, A. CONICS integrates scRNA-seq with DNA sequencing to map gene expression to tumor sub-clones. Bioinformatics 34, 3217–3219 (2018).

56. Gavish, A. et al. Hallmarks of transcriptional intratumour heterogeneity across a thousand tumours. Nature 618, 598–606 (2023).

57. Tirosh, I. et al. Dissecting the multicellular ecosystem of metastatic melanoma by single-cell RNA-seq. Science 352, 189–196 (2016).

58. Subramanian, A. et al. Gene set enrichment analysis: a knowledge-based approach for interpreting genome-wide expression profiles. Proc. Natl. Acad. Sci. U. S. A. 102, 15545–15550 (2005).

59. Newman, A. M. et al. Determining cell type abundance and expression from bulk tissues with digital cytometry. Nat. Biotechnol. 37, 773–782 (2019).

60. Wei, R. et al. Spatial charting of single cell transcriptomes in tissues. Nature biotechnology 40, 1190 (2022).

61. Fast Nearest Neighbor Search Algorithms and Applications [R package FNN version 1.1.4.1]. (2024).

62. Raredon, M. S. B. et al. Comprehensive visualization of cell-cell interactions in single-cell and spatial transcriptomics with NICHES. Bioinformatics 39, btac775 (2023).

